# Estimating effective population size from temporal allele frequency changes in experimental evolution

**DOI:** 10.1101/051854

**Authors:** Ágnes Jónás, Thomas Taus, Carolin Kosiol, Christian Schlötterer, Andreas Futschik

## Abstract

The effective population size (*N*_*e*_) is a major factor determining allele frequency changes in natural and experimental populations. Temporal methods provide a powerful and simple approach to estimate short-term *N*_*e*_. They use allele frequency shifts between temporal samples to calculate the standardized variance, which is directly related to *N*_*e*_. Here we focus on experimental evolution studies that often rely on repeated sequencing of samples in pools (Pool-Seq). Pool-Seq is cost-effective and outperforms individual-based sequencing in estimating allele frequencies, but it is associated with atypical sampling properties: additional to sampling individuals, sequencing DNA in pools leads to a second round of sampling increasing the estimated allele frequency variance. We propose a new estimator of *N*_*e*_, which relies on allele frequency changes in temporal data and corrects for the variance in both sampling steps. In simulations, we obtain accurate *N*_*e*_ estimates, as long as the drift variance is not too small compared to the sampling and sequencing variance. In addition to genome-wide *N*_*e*_ estimates, we extend our method using a recursive partitioning approach to estimate *N*_*e*_ locally along the chromosome. Since type I error is accounted for, our method permits the identification of genomic regions that differ significantly in *N*_*e*_. We present an application to Pool-Seq data from experimental evolution with *Drosophila*, and provide recommendations for whole-genome data. The estimator is computationally efficient and available as an R-package at https://github.com/ThomasTaus/Nest.

## INTRODUCTION

During experimental evolution studies, populations are maintained for many generations under specific laboratory conditions (Kawecki *et al.* 2012; SchlÖtterer *et al.* 2015; Long *et al.* 2015). For sexual organisms, such as *Drosophila*, the founder population is derived from a natural population and maintained in a new environment, typically at a census size of less than 2000 individuals. With such small population sizes genetic drift causes stochastic fluctuations in allele frequencies. Under neutrality, the level of random frequency changes is determined by the effective population size (*N*_*e*_) (Wright 1931). Furthermore, under certain conditions, the efficacy of deterministic forces, such as selection, is controlled by the population size (Charlesworth 2009). For weakly selected alleles the probability of fixation is directly proportional to the product of *N*_*e*_ and intensity of selection (Fisher 1930; Kimura 1964; Kimura 1964). Since changes in allele frequency are greatly affected by the population size, it is fundamental to estimate *N*_*e*_ in order to understand molecular variation of experimental evolution studies.

Krimbas and Tsakas (1971) estimated *N*_*e*_ using the standardized variance of allele frequency (*F*) from longitudinal samples in natural populations of olive flies. The underlying idea is that under neutral Wright-Fisher evolution, the variance of allele frequency after *t* generations is well described by the starting allele frequency and the drift level, which is inversely proportional to *N*_*e*_ (see also Falconer and Mackay (1996)). However, the frequencies are estimated from a sample, such that *F* must be corrected for the random errors due to sampling from the entire population. Nei and Tajima (1981) derived *N*_*e*_ estimates under two different sampling plans. Nei and Tajima (1981) and Pollak (1983) developed improved measures of *F* with reduced variance. Finally, Waples (1989) introduced a generalized framework that unified the different sampling plans introduced by Nei and Tajima (1981). Using their framework Jorde and Ryman (2007) proposed an alternative estimator of *N*_*e*_ that weights alleles differently and reduces the bias of the previous estimators.

With the advent of powerful computers, maximum likelihood-based methods became increasingly popular (Williamson and Slatkin 1999; Anderson *et al.* 2000; Wang 2001; Hui and Burt 2015) in addition to the moment-based approaches discussed above. Likelihood methods have the advantage that they can be extended beyond examining neutrally evolving isolated populations. For example, they permit to include migration (Wang and Whitlock 2003) and selection (Bollback *et al.* 2008; Malaspinas *et al.* 2012; Mathieson and McVean 2013). Although, these methods show less bias than the moment-based approaches (Wang 2001), they are still computationally demanding, in particular for the large numbers of markers typically obtained with novel sequencing technologies (Foll *et al.* 2015).

Estimating *N*_*e*_ with temporal methods requires samples collected at least at two time points. Alternative methods that use only a single time point are based on linkage disequilibrium (LD) (Hill 1981; Waples and Do 2008; Waples and Do 2010; Waples and England 2011), heterozygote excess (Pudovkin *et al.* 1996), molecular co-ancestry (Nomura 2008), sibship frequencies (Wang 2009; Wang 2013) or combination of summary statistics using approximate Bayesian computation (Tallmon *et al.* 2008). LD-based methods require haplotype data, which limits their applicability to large-scale population studies.

Although the cost for sequencing has dropped considerably, the separate sequencing of thousands of individuals in replicate populations in experimental evolution studies is still out of reach. Sequencing samples in pools (Pool-Seq) can provide a cost-effective alternative (SchlÖtterer *et al.* 2014). Pool-Seq has also been shown to outperform individual-based sequencing in estimating allele frequencies and inferring population genetic parameters under several conditions (Futschik and SchlÖtterer 2010; Zhu *et al.* 2012; Gautier *et al.* 2013). For these reasons Pool-Seq has become the basis of many experimental evolution ‘Evolve and Resequence’ (E&R) studies (Turner *et al.* 2011; SchlÖtterer *et al.* 2015). Following the emergence of E&R, many population genetic estimators have been adjusted to handle the properties of Pool-Seq data (Futschik and SchlÖtterer 2010; Kofler *et al.* 2011; Kofler *et al.* 2011; Kolaczkowski *et al.* 2011; Ferretti *et al.* 2013; Boitard *et al.* 2013). To the best of our knowledge, no *N*_*e*_ estimators have been used so far which account for the peculiarities of Pool-Seq.

In this article, we present a novel temporal method to estimate *N*_*e*_ from pooled samples. We show that previously proposed estimators can lead to substantial biases, as they neglect the variance component due to pooled sequencing. We introduce a new model accounting for the two stage sampling process associated with Pool-Seq data. In the first sampling step individuals are drawn from the population to create pooled DNA samples. In the second step, pooled sequencing is modeled as binomial sampling of reads out of the DNA pool. We show on simulated data that our method outperforms classical temporal *N*_*e*_ estimators. We then extend our estimators to heterogeneous effective population sizes along a chromosome. Partitioning DNA sequences into a number of segments is used to obtain separate *N*_*e*_ estimates for each segment. Finally, we present an application to a genome-wide experimental evolution data set from *Drosophila melanogaster* (Franssen *et al.* 2015).

## MATERIALS AND METHODS

### Two sampling schemes

Nei and Tajima (1981) pointed out that our ability to accurately estimate the population size depends on assumptions about the method of sampling individuals for genetic analyses. They proposed two different sampling schemes. Under the first scheme (plan I), individuals are either sampled after reproduction, or returned to the population after examining their genotypes. Nei and Tajima (1981) used the hypergeometric distribution to model draws of individuals from a starting population. This restricts the analysis to the assumption that actual size of the population is equal to the effective size. In contrast, under the second scheme (plan II) sampling takes place before reproduction and the sampled individuals are permanently removed form the population and their genotypes do not contribute to the next generation. The authors applied plan II assuming that the ratio between census and effective population size is large, so that sampling does not have an effect on the underlying allele frequency dynamics. In such situations binomial sampling can be used as an approximation.

Waples (1989) considered binomial sampling out of an infinitely large parental gamete pool for both sampling schemes. He concluded that the measure of variance under the two sampling plans differs only in a covariance term. For plan I, there is a positive correlation between allele frequencies sampled *t* generations apart because they are both derived from the same population at generation 0. In contrast, for plan II, the initial sample and individuals contributing to the next generation can be considered as independent binomial samples, thus sample frequencies at generation 0 and *t* are uncorrelated.

Contrary to the previous approaches, we consider two distinct sampling steps at each time point (Fig. 1). In the first step, we sample individuals out of the population to create pooled samples for sequencing. Sampling individuals can take place according to either plan I or plan II. In the second step, we model drawing reads out of the pooled DNA sample. The estimated allele frequency variance is then corrected for the additional variance coming from both sampling steps.

**Figure 1:**
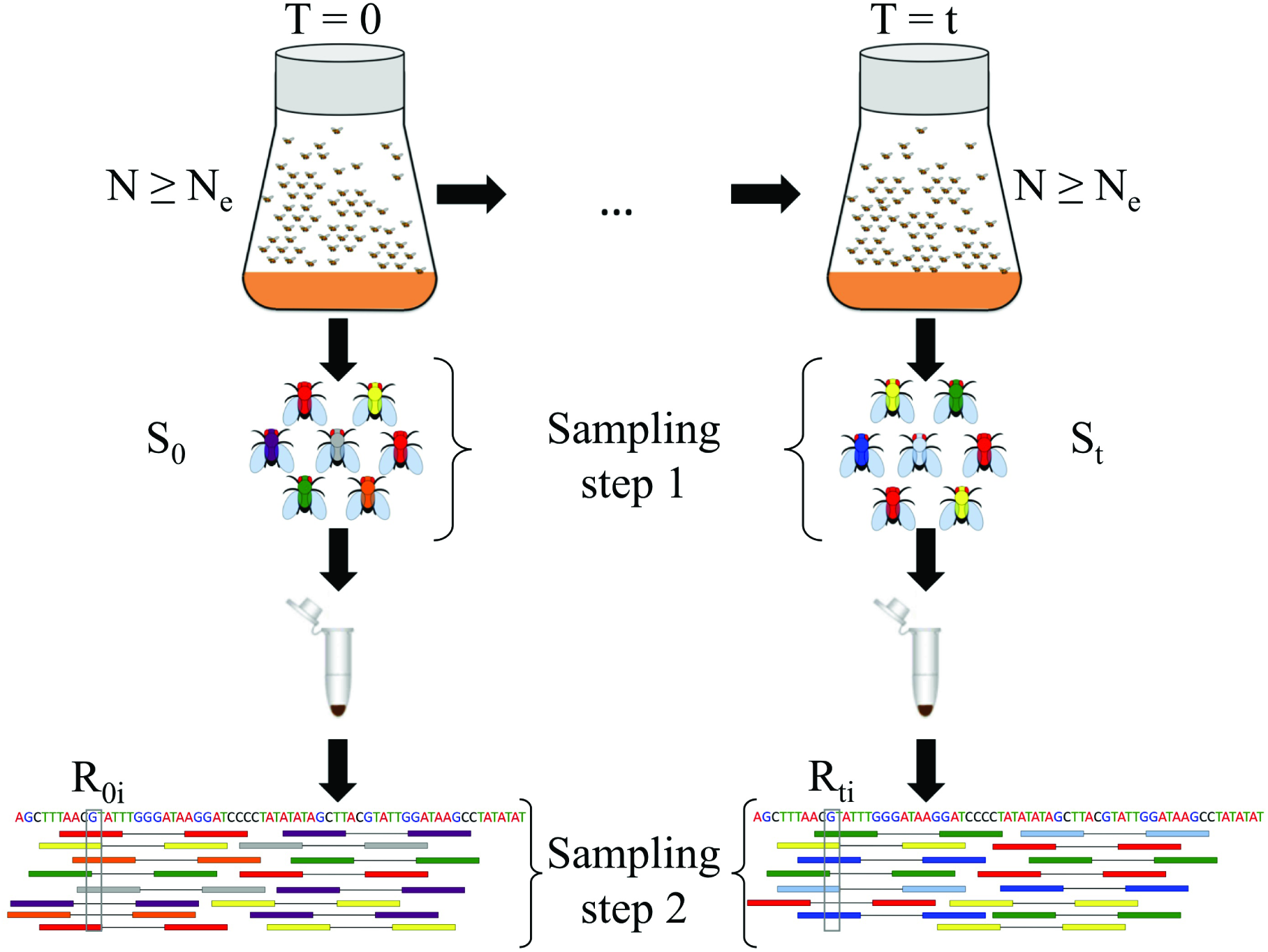
Two-step sampling in experimental evolution with *Drosophila*. In E&R studies, populations are propagated at a census size (*N*) defined by the experimenter, which is, in general, larger than the effective population size *N*_*e*_. Using temporal methods *N*_*e*_ can be estimated from the variance in allele frequency in samples taken *t* generations apart. To get an accurate representation of allele frequencies in population genetic studies a large number of individuals *S*_*j*_ (*j* ∈ {0, *t*}) are sampled and pooled. Sampling can take place according to sampling plan I or II based on the mode of reproduction. Pooled samples are then subjected to high throughput sequencing. We represent random variation in sequence coverage (*R*_*j*_, *j* ∈ {0, *t*}) with an additional sampling step (called sampling step 2). We correct for both sampling steps when estimating *N*_*e*_ in pooled samples. Additionally, we take into account variable coverage levels across the genome (coverage *R*_*ij*_ for site *i* at *T* = *j*) when correcting for the variance coming from sequencing.

For a typical E&R study, outbred experimental populations are created by mixing a large number of inbred lines (Turner and Miller 2012; Bastide *et al.* 2013; Huang *et al.* 2014; Franssen *et al.* 2015). The populations are then propagated under the desired experimental conditions while keeping the census size of the population (*N*) controlled through time (Fig. 1). However, the experimenter has no direct influence on the effective populations size, which is in general, lower than the census size. In E&R studies with *Drosophila* the census size rarely exceeds some hundreds of individuals, and sampling usually takes place after reproduction according to plan I. For organisms maintained at larger sizes, such as yeast, the sample for genetic analysis is not returned to the population (Burke *et al.* 2014). Plan II applies to such cases. Sampled individuals are subsequently pooled together. The size of the pool can be as large as the whole population. Depending on the experimental design, it is also possible that only a fraction of the population is sequenced, for instance, only females (Tobler *et al.* 2014; Franssen *et al.* 2015). Pooled individuals are used to create DNA libraries, which are in turn, subjected to high throughput sequencing. Sequenced reads are mapped to a reference genome and filtered for subsequent analyses. Randomness in sequencing and local structures in the genome lead to a variable coverage level, which we also take into account when estimating *N*_*e*_.

### Notation

We use observed allele frequencies sampled *t* generations apart to estimate *N*_*e*_ (Fig. 1). We assume that the population is propagated at constant census size *N*, and that *N* ≥ *N*_*e*_. We consider only biallelic sites. At each locus the true population allele frequency at time *T* = *j* is denoted by *p*_*j*_, where *j* ∈ {0, *t*}. To obtain allele frequency estimates for the unknown *p*_*j*_, the population is subjected to sampling. In contrast to previous approaches, we consider two sampling steps (Fig. 1). At *T* = *j*, we first sample *S*_*j*_ individuals out of the population to create a pooled sample for sequencing. Sampling can take place according to either plan I or plan II, as described above (also shown in Fig. S1). For the second sampling step, we model Pool-Seq by drawing *R*_*ij*_ reads out of the pooled DNA sample at site *i*. This permits for variation in sequence coverage. Below we denote the unknown sample allele frequency among the *S*_0_ individuals at the first sampling time point (*T* = 0) by *x*, and the subsequent allele frequency estimate obtained via pool sequencing from *R*_0_ reads by 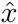. Similarly, at some *T* = *t*, the respective frequencies are denoted by *y* and *ŷ*. Note that under pool sequencing only 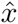 and *ŷ* are observed.

### Estimating N_e_ from temporal allele frequency changes

Under neutral Wright-Fisher evolution the variance in allele frequency 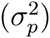 generated by drift after *t* generations at a single locus in a diploid population is well described by the following expression

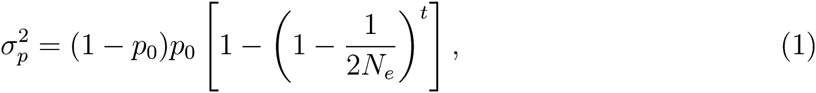

where _*p*0_ is the starting allele frequency (Falconer and Mackay 1996). Wright (1931) denoted the standardized variance by *F* = 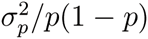, which leads to a convenient closed form expression for *N*_*e*_. Furthermore, if *N*_*e*_ is large enough, *F* ≈ 1 — *e^−t^/^2*N*_*e*_^* and *N*_*e*_ can be calculated as

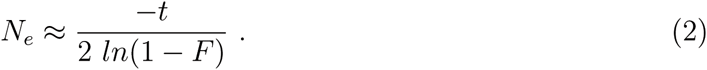

The relation between *N*_*e*_ and allele frequency changes described in equation (1) was first used by Krimbas and Tsakas (1971) in natural populations of olive flies. They estimated the variance using

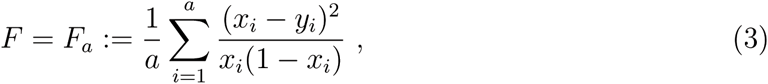

where *x*_*i*_ and *y*_*i*_ (*i* = 1, …, *a*) are the observed allele frequencies in the samples collected *t* generations apart and a is the number of alleles at *a* specific locus. To eliminate the contribution of sampling errors to the variance, the total variance *F*_*a*_ was corrected for the random sampling noise by simply subtracting the corresponding binomial variance. This approach was further investigated and developed by a number of authors (Pamilo and Varvio-Aho 1980; Nei and Tajima 1981; Pollak 1983; Waples 1989).

Possible sources of bias in *N*_*e*_ estimators were later investigated by Jorde and Ryman (2007). The authors pointed out that the expectation over *F* is typically approximated by taking the expected values separately for the numerator and the denominator (Turner *et al.* 2001). They suggested a different weighting scheme of alleles leading to an alternative unbiased estimator to measure temporal frequency change.

### Correction for two stage sampling

We follow the sampling approach described above, when calculating the correction term for pooled samples. We consider a diploid, random mating population of size N with discrete generations. Neutral evolution is assumed with no selection, migration and mutation. Samples are drawn from the population at generation *T* = 0 and *t*. To model sampling variation, we assume two stage sampling for both sampling plans as illustrated in Fig. S1. In the first step, we model drawing *S*_*j*_ individuals out of *N* to create pooled sequencing samples. We consider diploid populations throughout the derivation, so that a sample of *S*_*j*_ individuals leads to 2*S*_*j*_ sequences. Sampling is assumed to be binomial with parameters 2*S*_*j*_ and *p*_*j*_ (Waples 1989). In the second step, pool sequencing a random number *R*_*j*_ of reads is also modeled as binomial sampling.

By calculating the variance of the observed allele frequencies in pooled samples, we are able to correct for the variation introduced by the two-step sampling. Following Jorde and Ryman (2007), we use the following expression as the measure for the temporal change in allele frequency

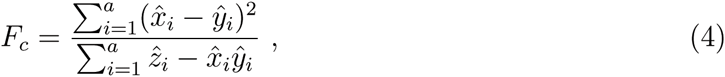

where *ẑ*_*i*_ = 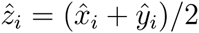 and *a* is the number of alleles (Nei and Tajima 1981). We consider only biallelic sites, which reduces *F*_*c*_ to 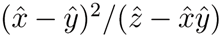 for a locus. The expectation of *F*_*c*_ is approximated in the following way

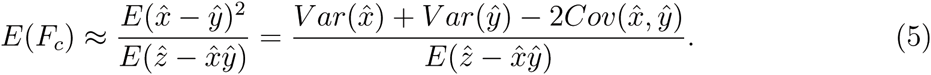

For both plans, we derive expressions for the numerator and denominator in equation (5) separately. Here we summarize our main conclusions, details on the derivation are provided in the supplementary material. With 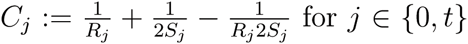, and *p* denoting the true population allele frequency in the gamete pool at generation 0, we obtain

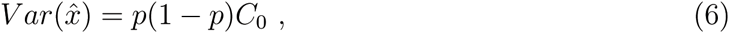

and

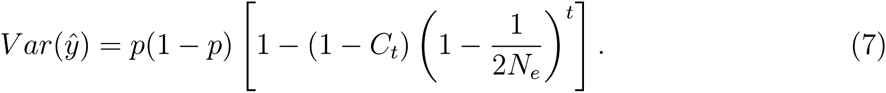

Following Waples (1989) the denominator in equation (5) reduces to

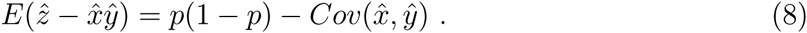

For plan II, the sample allele frequencies at generation 0 and *t* are uncorrelated, 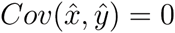 and *F*_*c*_ corrected for the noise coming from the two stage sampling is given by

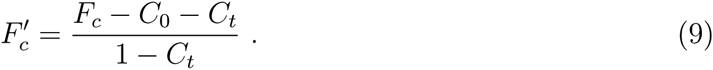

For plan I, on the other hand, the sample allele frequency at generation 0 is positively correlated to the sample allele frequency at *t* because both are derived from the same population at generation 0. This requires to calculate the sample covariance 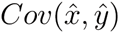 in equation (5). It turns out (see supplementary material for details) that the covariance of 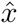 and *ŷ* is equal to the variance of *p*_0_

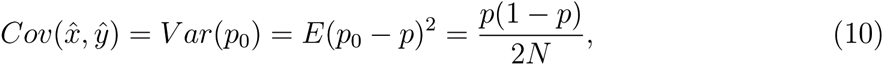

where *N* is the census size of the population at generation 0. Substituting the inferred covariance into equation (5) leads to the following corrected variance estimate, *F’*_*c*_ for plan I

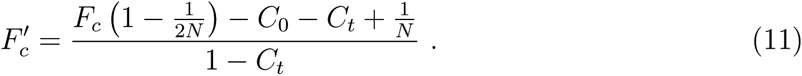

*N*_*e*_ is then estimated using the corrected variance estimates (equations (9) and (11)) instead of *F* in equation (2).

Having long time series, often spanning hundreds of generations, has recently become commonplace (Barrick *et al.* 2009; Burke *et al.* 2010; Burke *et al.* 2014). The assumption of *t* being small, often made when calculating *N*_*e*_ (Krimbas and Tsakas 1971; Nei and Tajima 1981), can lead to severe bias when approximating 2*N*_*e*_ ≈ *t*/*F*. Therefore, instead of assuming *t* being small, we use the formula in equation (2) for estimating *N*_*e*_.

In our genome-wide data set, we apply individual correction terms *C*_*ij*_ (*j* ∈ {0,*t*}) for each marker *i* (*i* = 1,… *n*) before averaging over the *n* biallelic loci in a window.

### Simulations

We evaluate the performance of our novel estimator on simulated data. Using the neutral Wright-Fisher model, we simulate allele frequency trajectories at *n* independent loci (SNPs) in a population of *N*_*e*_ diploid individuals. Usually our simulations start with uniformly distributed allele frequencies. Nevertheless, we also assess the performance of our estimator on simulated data with a starting allele frequency spectrum skewed towards low frequency variants resembling neutral expectations. To investigate the effect of the ratio between census and effective population size (*r* = *N*/*N*_*e*_), we increment the population size to a desired level of *N* while keeping the allele frequencies unchanged before sampling takes place. At the start and after *t* generations samples are taken out of the population. Pooling individuals is modeled by sampling without replacement. Genome-wide sequence data typically has uneven coverage among sites, which we model by a Poisson distribution with a given mean coverage. For every biallelic genomic position, the number of reads corresponding to the target coverage is determined by binomial sampling. *N*_*e*_ is estimated between the start and after *t* generations for *n* =1000 SNPs in 100 replicate simulations. We report summary statistics of the resulting *N*_*e*_ estimates across replicates.

Linkage disequilibrium between loci can reduce the number of independent SNPs, thereby increasing the variance of the estimate. The impact of non-independence between SNPs is evaluated using whole-genome forward simulations with recombination performed by the software tool MimicrEE (Kofler and SchlÖtterer 2014). We generate a diploid founder population consisting of 2000 simulated haplotypes that mimic a wild population of *D. melanogaster* from Vienna (Bastide *et al.* 2013; Kofler and SchlÖtterer 2014) and use the recombination rate of *D. melanogaster* (Fiston-Lavier *et al.* 2010). Forward simulations are performed in 10 replicate runs. Allele counts are subjected to binomial sampling to mimic Pool-Seq with a given sequence coverage. *N*_*e*_ is estimated for non-overlapping windows, each containing a fixed number of SNPs.

### Estimating N_e_ on simulated data

We denote our estimator corrected for the additional sampling step, i.e. pooling, by *N*_*e*_(*P*). We compare the performance of *N*_*e*_(*P*) to methods that correct only for a single sampling step. We assume binomial sampling out of an infinite gamete pool recommended by Waples (1989) and the weighting scheme of alleles put forward in Jorde and Ryman (2007). Therefore we decided to compare *N*_*e*_(*P*) to the estimators proposed by these authors, and denote them by *N*_*e*_(*W*) and *N*_*e*_(*JR*), respectively.

We illustrate experimental sampling procedures by considering two major scenarios: (i) the full population is sequenced as one large pool; (ii) only a subset of the population is used to create pooled samples. Under scenario (i) we perform only a single binomial sampling step to represent sampling reads out of the DNA pool. The pool size is set to be equal to the census size of the population (*S*_*j*_ = *N*) for our estimators. In this case, the coverage represents the sample size for estimators that correct only for a single sampling step. For scenario (ii), we sample individuals without replacement to generate pools of *S*_*j*_ individuals that are subjected to sequencing. Sequencing reads is modeled by binomial sampling as before. For this scenario again the coverage is taken as the sample size for *N*_*e*_(*W*) and *N*_*e*_(*JR*) estimators. Note, however, that the latter estimators permit correction only for a single sampling step, therefore the user has to decide on whether sampling individuals or reads is corrected for. In general, it is recommended to correct for the sampling step that contributes a larger extent to the sampling variance.

Genome-wide coverage differences are treated equally by subtracting correction defined by the corresponding coverage for each site before averaging over the number of SNPs for all methods.

### Change point inference for the genome-wide estimates

Selection causes change in allele frequency at a targeted site and also affects nearby sites through linkage. Consequently, deviation from the neutral expectation can result in a locally decreased *N*_*e*_. To detect patterns of heterogeneous *N*_*e*_ along the genome we partition chromosomes into windows of locally homogeneous *N*_*e*_. To distinguish random fluctuations from systematic changes, we applied a segmentation algorithm, which guarantees type I error control, in the sense that the estimated number of windows will not exceed the true one except for a small error probability *α* to be specified by the user. The algorithm is implemented in the stepR software package (Frick *et al.* 2014). It is related to a statistical multiscale change-point estimator (SMUCE) that has been suggested for partitioning sequences with respect to GC-content in (Futschik *et al.* 2014). As this method requires homogeneous variances, we applied it to log-transformed *N*_*e*_ estimates calculated genome-wide for non-overlapping windows. For visualization we back-transformed the obtained step function.

## RESULTS AND DISCUSSION

### Two-step correction is vital to avoid large bias in *N*_*e*_ estimates for long time series

Methods that do not correct for the additional sampling step, i.e. pooling, can lead to substantial bias in *N*_*e*_ estimates. We illustrate this (Fig. 2) on simulated data where the effective population size is inferred using allele frequency samples from the initial and evolved populations. We compare our proposed estimator (*N*_*e*_(*P*)) to two commonly used estimators *N*_*e*_(*W*) (Waples 1989) and *N*_*e*_(*JR*) (Jorde and Ryman 2007). Figure 2 shows that the additional correction substantially decreases the bias for almost all data points. Methods that account only for a single sampling step may be severely biased. For the estimator *N*_*e*_(*JR*) the bias becomes less pronounced with an increasing number of generations. Under plan I, *N*_*e*_(*P*) is nearly unbiased, and plan II has a slight upward bias when applied on data simulated under plan I and the samples are taken at very close time points. When samples are collected only a few generations apart, the variance of *N*_*e*_(*P*) estimators tends to be larger than that of *N*_*e*_(*W*) and *N*_*e*_(*JR*) under both plans.

**Figure 2:**
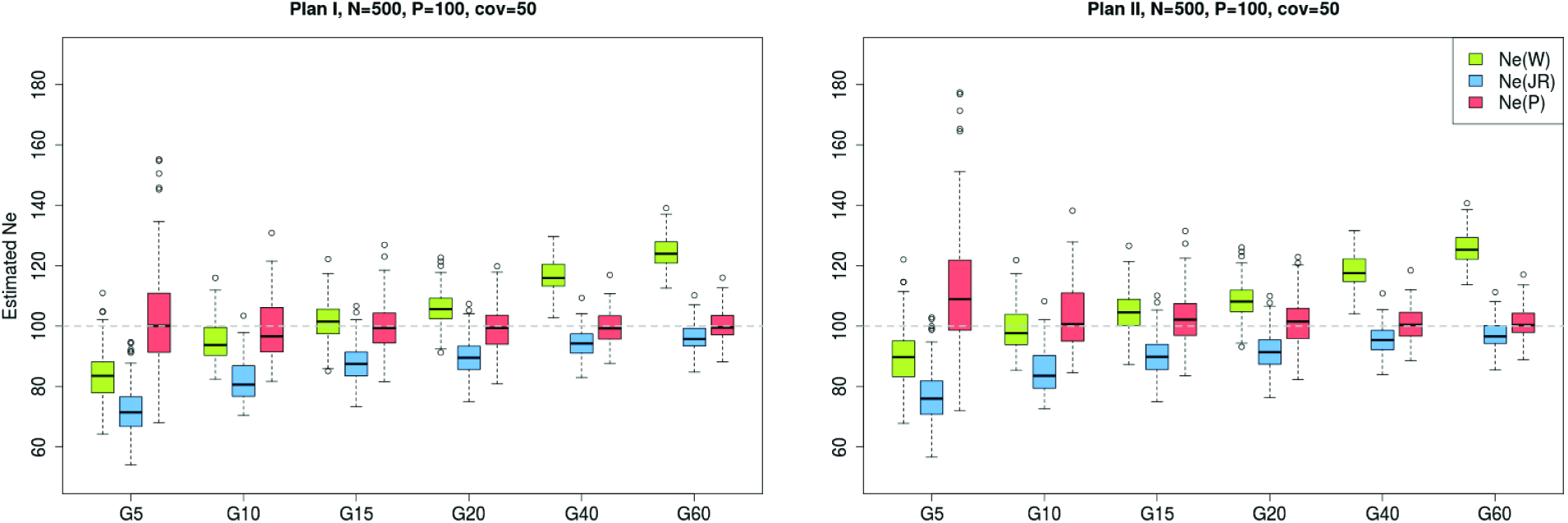
Performance of different methods to estimate *N*_*e*_ based on simulated data. Sixty generations of Wright-Fisher neutral evolution with *N*_*e*_ = 100 diploid individuals were simulated for 1000 unlinked loci (SNPs). Prior to sampling, the population was increased to a census size of N = 500 individuals at each generation. At the starting population (G0), and at each indicated time point a sample was taken to create a pool of P=100 individuals. Sequencing was modeled with an additional subsequent binomial sampling step of a random number of reads (50 on average) for each SNP. Then *N*_*e*_ was estimated on the resulting data set by separately contrasting G0 with each of the generations G5, G10, G15, G20, G40 and G60. We considered the estimators *N*_*e*_(*W*) (Waples 1989), *N*_*e*_(*JR*) (Jorde and Ryman 2007) and our *N*_*e*_(*P*). Each box represents results from 100 simulations with identical parameters. The dashed grey line shows the true value of *N*_*e*_. Data is simulated under plan I assumptions and the results of plan I and II estimators are shown in the left and right panel, respectively.

Plan I and II estimators differ by a factor resulting from the covariance between the sample frequencies at generation 0 and *t* (equation (10)), which is inversely proportional to the census population size. Consequently, the difference between plan I and II becomes smaller for increasing *N*. Waples (1989) investigated how the ratio between census and effective population size (*r* = *N*/*N*_*e*_) affects the accuracy of the estimators, and concluded that the ratio of *r* ≥ 2 is sufficient to reach similar estimates for both sampling schemes. We tested the performance of *N*_*e*_(*P*) on simulated data with *N*_*e*_ = 100 and *N*: *N*_*e*_ ratios of *r* = 1,2, 5 with different coverages and pool sizes (Fig. S2-S4). When *N* = *N*_*e*_, the *N*_*e*_(*P*) plan I method achieves highly accurate estimates for all time points in contrast to the other methods (Fig. S2). If, however, *N*_*e*_(*P*) plan II estimator is applied to data simulated under plan I, we observe an upward bias for small *t*, which improves with an increasing number of generations. This pattern is not unexpected since the missing covariance term becomes less influential in view of the increasing drift variance with more generations. When the entire population is sequenced as a single pool (*P* = 100), the plan II estimators of Waples (1989) and Jorde and Ryman (2007) perform similarly to the *N*_*e*_(*P*) plan I estimator because the correction for pooling in *N*_*e*_(*P*) cancels out the additional covariance term when *P* = *N* making the term used as F approximately identical to that of *N*_*e*_(*JR*). This is a general pattern irrespective of *r*.

For *r* ≥ 2, *N*_*e*_(*P*) plan I remains highly accurate (Fig. S3 and S4). Furthermore, when increasing the census size under a constant *N*_*e*_ (equivalent to increasing *r*), the covariance between sample allele frequencies decreases making the difference between plan I and II almost negligible (Waples 1989). Also, the sampling variance becomes proportionally smaller compared to the drift variance with increasing number of generations between the samples. This improves our ability to accurately estimate *N*_*e*_.

Correcting for the additional variance inherent to Pool-Seq leads to an improved performance of *N*_*e*_(*P*) compared to the classical methods for both plans. In general, with Pool-Seq data the extent of the bias in *N*_*e*_(*W*) and *N*_*e*_(*JR*) estimates depends on the ratio between *N* and *P*, smaller pool sizes leading to a larger bias. Since we accounted for the sequencing step with these estimators (see section Estimating *N*_*e*_ on simulated data), decreasing the coverage at a given pool size does not change the bias to a large extent but rather increases the variance of the estimators.

In most of the experimental studies the investigator has control over the census population size, thus requiring the knowledge of *N* for *N*_*e*_(*P*) plan I does not restrict the analysis. We illustrate the performance of *N*_*e*_(*P*) plan I only in subsequent simulations where *N*_*e*_ = *N* is used but according to Fig. S3 and S4 *N*_*e*_(*P*) plan I is also highly accurate when *r* ≥ 2.

We show the coefficient of variation (CV) of the *N*_*e*_(*P*) plan I estimator in Fig. 3. CV is defined as the ratio between the standard deviation and the mean (*CV* = *σ*/*μ*, where both *σ* and *μ* are estimated from the sample). It measures the relative dispersion of the distribution of the estimated values. *N*_*e*_(*P*) estimators are highly precise in nearly all cases, except when the drift variance is negligible compared to the sampling variance (Fig. 3 see also Fig. S6 and S8 where *N*_*e*_ = 1000, *t* < 30, *P* ≤ 100 and *c* = 50). The bias is coming from a few oulier estimates, but the median shows more robust results (Fig. S10 and S11). For plan II estimators, the behavior of the method is similar (Fig. S5 and S7,S9). Note that the simulations underlying Fig. S5 and S7,S9 have been done under plan I sampling scheme.

**Figure 3:**
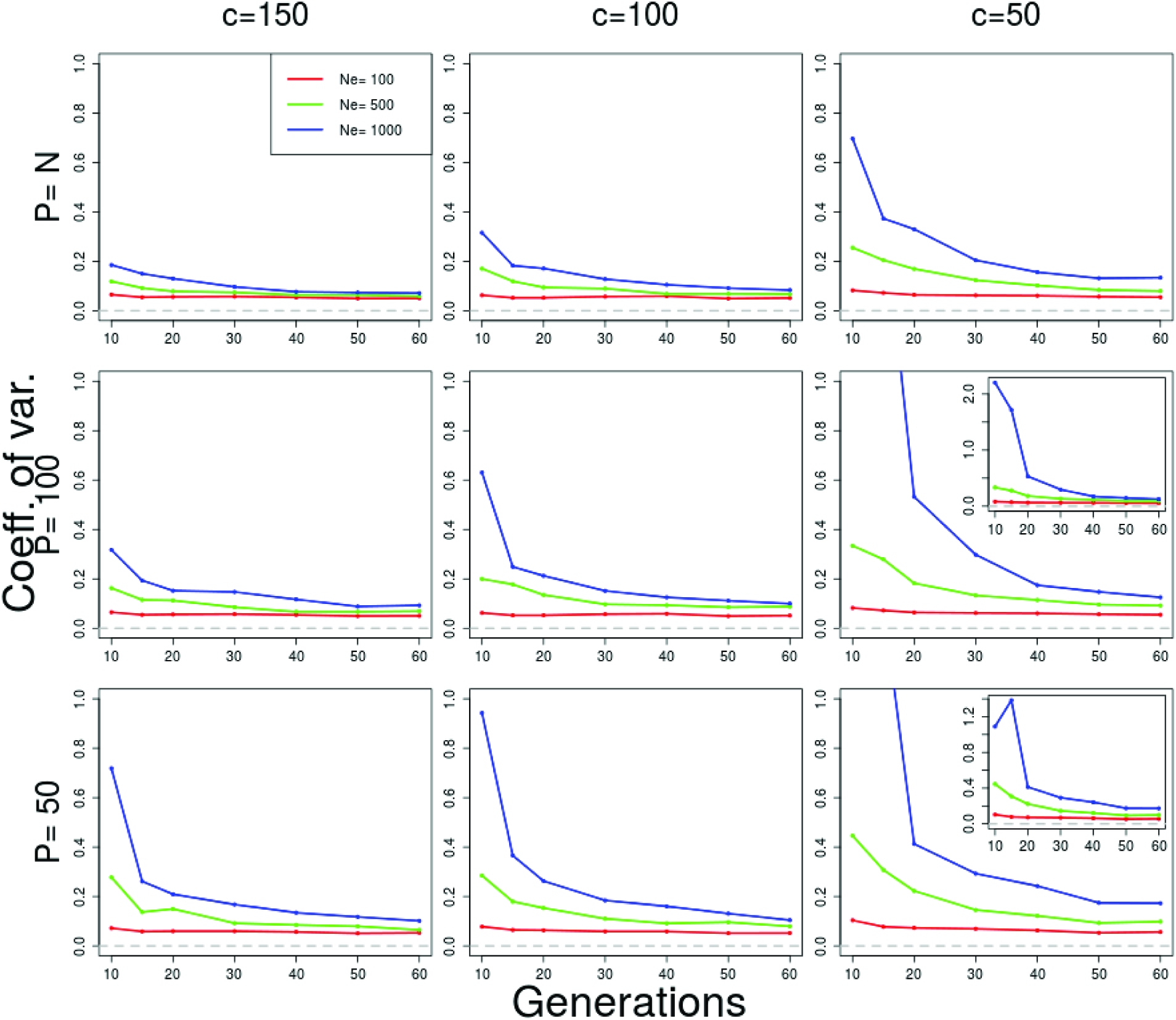
Coefficient of variation of *N*_*e*_(*P*) for plan I for various parameter values. Neutral Wright-Fisher simulations were performed with various combinations of the parameters effective population size (*N*_*e*_ = 100, 500, 1000 diploid individuals), pool size (*P* = 100, 50) and coverage (*c* = 150, 100, 50). P = N indicates scenarios when the whole population is sequenced as a single pool. For all simulations, *N* = *N*_*e*_ was used as census size of diploid individuals. Each value is calculated over 100 simulations. Negative estimates were excluded when calculating summary statistics. The frequency of negative estimates is listed in table S1. In situations where the coefficient of variation exceeds one, an inset figure shows the actual value.

### Increasing the number SNPs reduces the variance of N_e_(P)

We test how the number of loci used to infer *N*_*e*_ affects the accuracy and the precision of the estimates by gradually increasing the number of independent SNPs from 100 to 10000 (Fig. 4). We observe a larger variance and a slight downward bias for a small number of SNPs (100 SNPs). Both the bias and the variance become smaller with a larger the number of SNPs. Some further improvement is obtained when more than 10K SNPs are used (not shown), but the benefit of additional independent SNPs levels off. We conclude that *n* = 1000 independent SNPs usually provide sufficient precision and accuracy. However, when linkage disequilibrium is present in a genome-wide data set, the number of truly independent SNPs per window is reduced and a larger number of loci is recommended.

**Figure 4:**
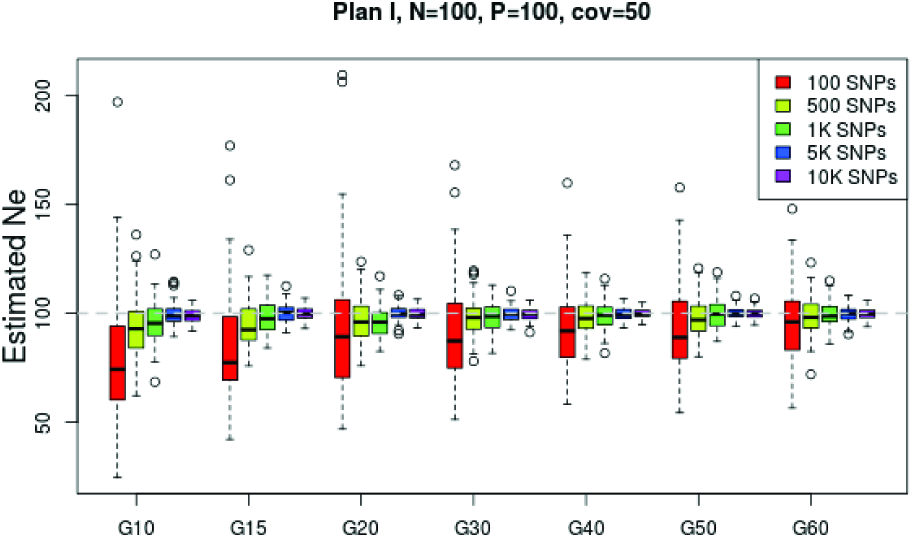
Effect of the number of SNPs used on the *N*_*e*_ estimates. The effective population size is estimated using *N*_*e*_(*P*) plan I on simulated data with *N*_*e*_ = *N* = 100. A total of *P* = 100 individuals is pooled and sequenced at a mean coverage of *c* = 50. Based on 100 simulation runs, *N*_*e*_ is estimated using different numbers of SNPs at multiple time points.

### Skew towards low frequency variants only moderately increases the variance of N_e_(P)

In natural populations, the neutral site frequency spectrum is skewed towards low frequency variants. *N*_*e*_(*P*) uses a weighting scheme that is not very sensitive to this skew, see also Jorde and Ryman (2007). This makes it robust with respect to the shape of the starting allele frequency distribution. We illustrate this with a simulated data set having a starting allele frequency distribution that is skewed towards low frequency variants (exponential) as predicted under neutrality. The estimates of *N*_*e*_ from such data sets are compared to simulated data with matching parameters but uniform starting allele frequency distribution (Fig. 5). We observe a very slight upward bias with neutral starting allele frequencies compared to uniform, and a moderate increase in the variance given *t* ≥ 15. With increasing number of generations the difference becomes negligible.

**Figure 5:**
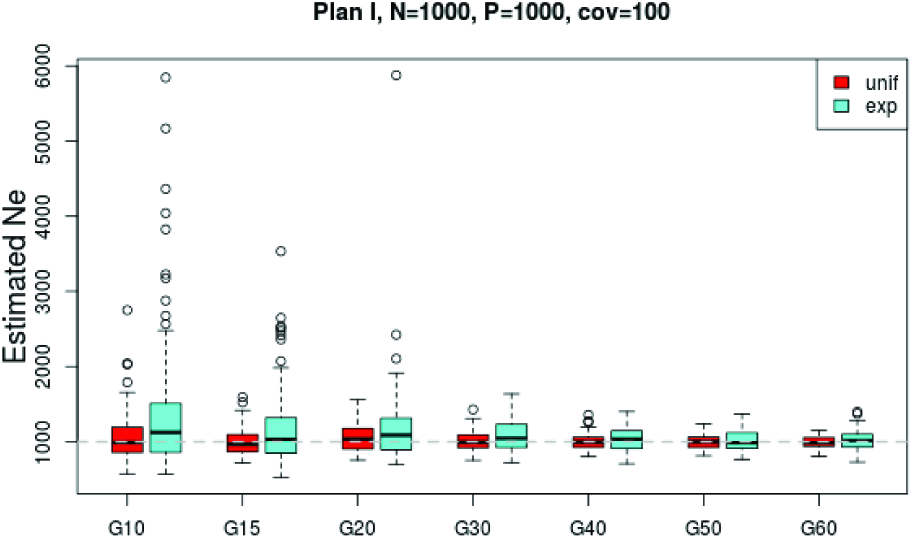
Influence of the starting allele frequency distribution on *N*_*e*_(*P*) plan I estimator. A comparison between uniform and exponentially distributed (neutral) starting allele frequencies is shown.

### The presence of linkage disequilibrium does not severely bias the estimate

We investigated the sensitivity of our estimator to linkage disequilibrium between loci using genome-wide neutral simulations with recombination (Kofler and SchlÖtterer 2014). To estimate the effect of linkage disequilibrium, we simulated three different rates of recombination: high, normal and no recombination. For the first case, the recombination rate is set to mimic the behavior of almost independent SNPs. In the normal recombination rate scenario, we use *D. melanoagster* recombination rates (Fiston-Lavier *et al.* 2010). The effective population size was estimated in non-overlapping windows with a fixed number of *n* = 5000 SNPs (Fig. 6). Different levels of linkage disequilibrium affect the number of independent loci per window. Nevertheless, we observe only a slight increase in the precision of the *N*_*e*_ estimates with increasing recombination rate (Fig. 6).

**Figure 6:**
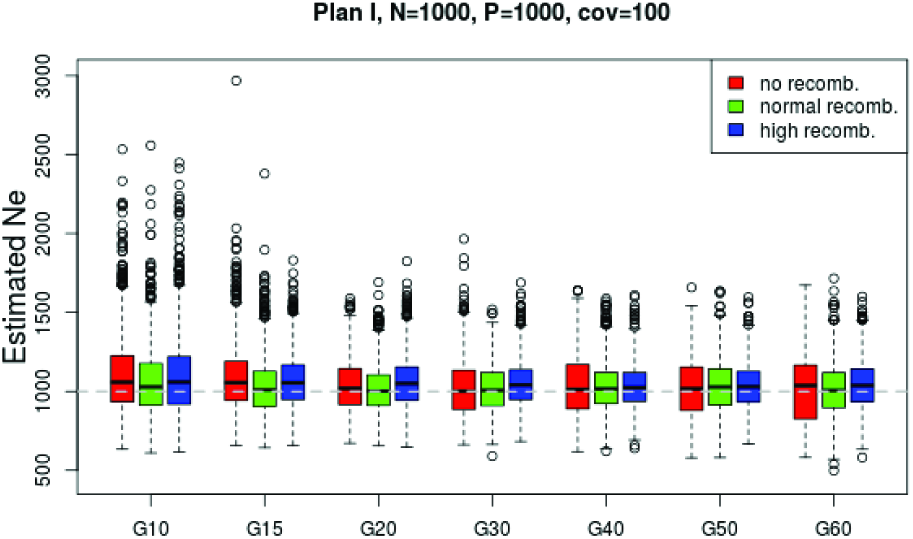
Effect of linkage disequilibrium. The effect of linkage disequilibrium on our estimators was evaluated based on a whole-genome forward simulation with recombination using the software MimicrEE (Kofler and SchlÖtterer 2014). Three sets of simulations were performed with different rates of recombination: high, normal and no recombination. For each parameter setup, a genome-wide simulation is replicated ten times. The effective population size was estimated with *N*_*e*_(*P*) plan I in non-overlapping windows of 5000 SNPs for each replicate. The distribution of *N*_*e*_ estimates across replicates and locations is shown in the box-plots.

### Genome-wide heterogeneity of N_e_ is recovered in real data of *D. melanogaster*

We analyzed data of a recent E&R study in *D. melanogaster* (Orozco-terWengel *et al.* 2012; Tobler *et al.* 2014; Franssen *et al.* 2015). In this experiment replicate populations of 1,000 individuals were subjected to a fluctuating hot environment for 59 generations. Allele frequency estimates were obtained for founder and evolved populations using Pool-Seq. *N*_*e*_ was estimated based on the allele frequency changes between founder and latest evolved populations using *N*_*e*_(*P*) under plan I. We considered non-overlapping windows of 10000 SNPs. To determine statistically significant changes in *N*_*e*_ along the genome we use the software provided by Frick *et al.* (2014). This method requires homogeneity of variances. Since, the variance of estimates obtained with *N*_*e*_(*P*) increases with *N*_*e*_, the estimates were log-transformed before applying the partitioning procedure. The obtained step function was back-transformed to the original scale and is shown for four biological replicates (Fig. 7).

**Figure 7:**
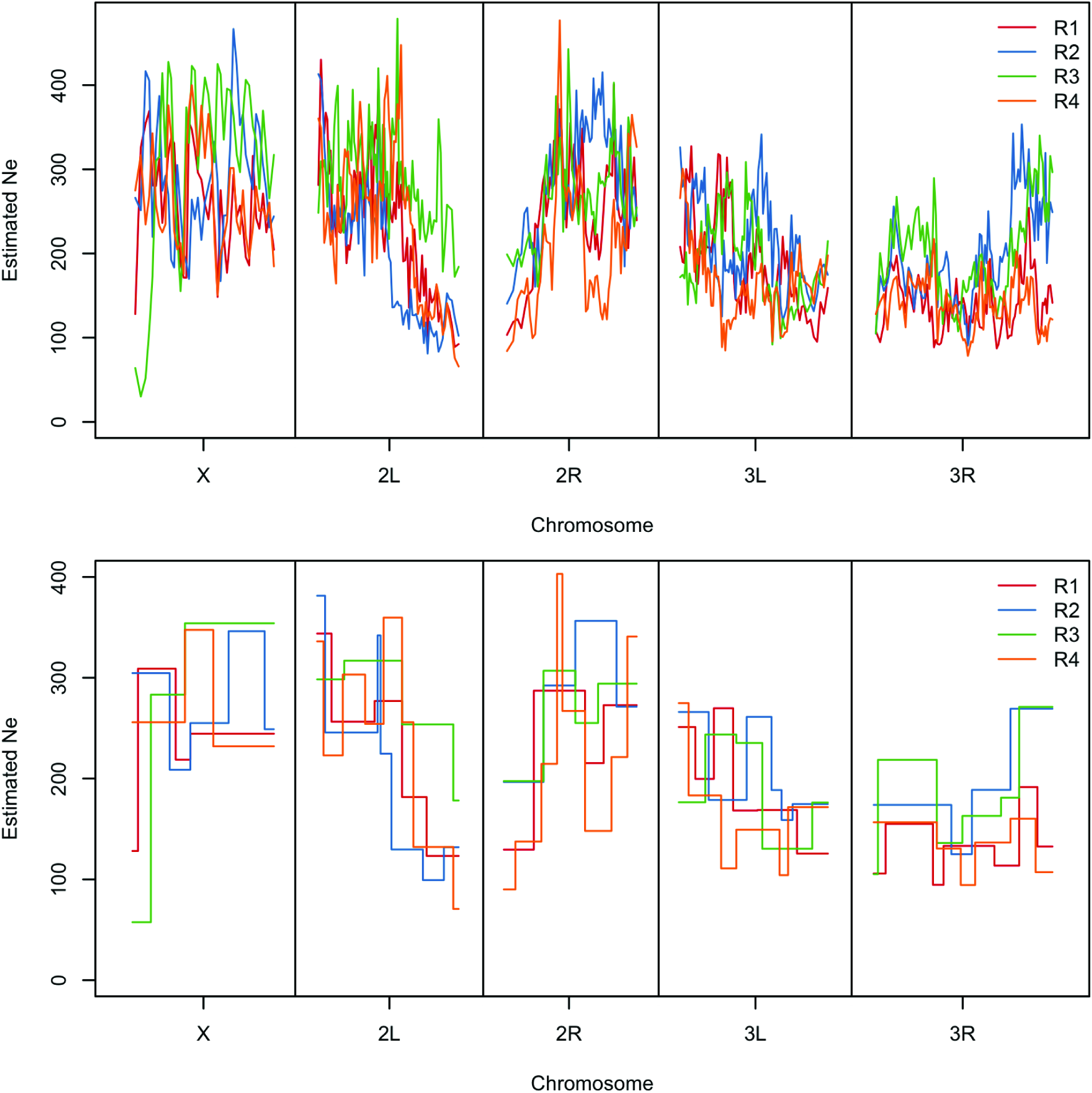
Genome-wide *N*_*e*_ estimates from an E&R study in *D. melanogaster. N*_*e*_ is estimated based on the allele frequency changes between founder and evolved populations at generation 59 (Franssen *et al.* 2015). In the top panel genome-wide estimates calculated with *N*_*e*_(*P*) plan I using non-overlapping windows of 10000 SNPs are shown. Chromosome-wide mean estimates are shown in table 1. Statistically significant changes in *N*_*e*_ is determined after log-transformation of the data (bottom panel). Chromosome arms are separated by vertical bars. Interestingly, the estimates on the X chromosome do not reflect the expectation of *N*_*e*_ values reduced by three-quarters compared to autosomes.

The mean estimates for each chromosome arm as well as across the genome is stable across replicates (see table 1). As experimental evolution studies often aim to find signals that are consistent across replicates, this can be an important check of the experimental set up. On the other hand, we see differences between chromosome arms. For example, the mean is clearly lower for 3R, emphasizing the added value of spatial analysis compared to genome-wide estimates.

**Table 1:**
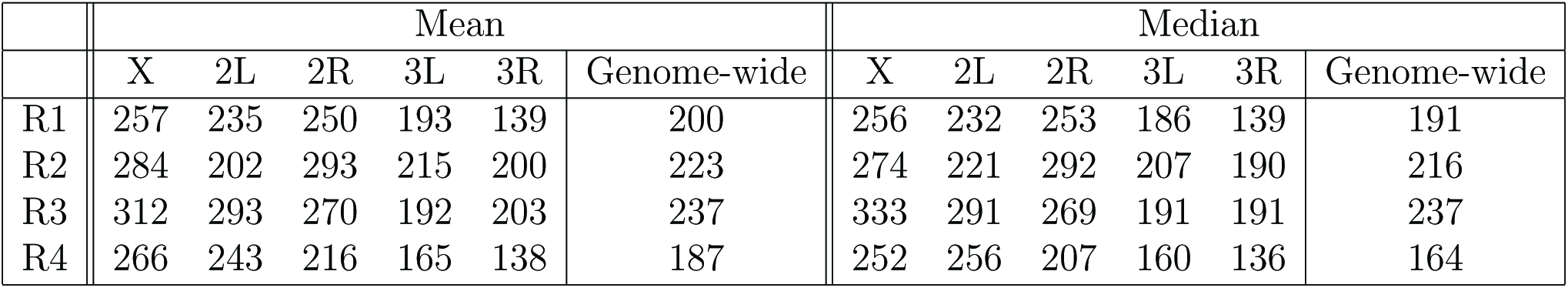
Genome-wide mean and median *N*_*e*_ estimates from an E&R study in *D. melanogaster*. The effective population size is estimated with *N*_*e*_(*P*) plan I in windows of 10000 SNPs (Fig. 7). The mean and median estimates across windows are shown for the major chromosome arms. Genome-wide mean and median is taken over the autosomes excluding chromosome 4.

|  | Mean |  |  |  |  |  | Median |  |  |  |  |  |
| --- | --- | --- | --- | --- | --- | --- | --- | --- | --- | --- | --- | --- |
|  | X | 2L | 2R | 3L | 3R | Genome-wide | X | 2L | 2R | 3L | 3R | Genome-wide |
| R1 | 257 | 235 | 250 | 193 | 139 | 200 | 256 | 232 | 253 | 186 | 139 | 191 |
| R2 | 284 | 202 | 293 | 215 | 200 | 223 | 274 | 221 | 292 | 207 | 190 | 216 |
| R3 | 312 | 293 | 270 | 192 | 203 | 237 | 333 | 291 | 269 | 191 | 191 | 237 |
| R4 | 266 | 243 | 216 | 165 | 138 | 187 | 252 | 256 | 207 | 160 | 136 | 164 |

All estimators mentioned above assume neutrality. Directional selection, however, leads to deterministic allele frequency changes that generate a larger deviation from the expected frequency than it would be due to random genetic drift alone. Consequently, estimates of *N*_*e*_ will be lower in the presence of directional selection. In the *D. melanogaster* example, *N*_*e*_ ranges from around 50 to almost 400. Around the centromere of chromosome 2, the estimated *N*_*e*_ decreases by two-third in replicates 1, 2 and 4. Even more striking, the estimated *N*_*e*_ is low on the entire chromosome arm 3R and also parts of 3L. Overall, these patterns can be attributed to strong LD, caused either by low recombination rates around the centromeres (Chan *et al.* 2012) or segregating inversions (Kapun *et al.* 2014) in combination with selection potentially on rare variants. These results are consistent with Tobler *et al.* (2014), who observed a massive amount of outlier SNPs around the centromere of chromosome 2 and on 3R. Strong LD, potentially due to selection on rare variants, could explain this inflated number of hitchhikers. Interestingly, certain regions of the genome show extensive differences in *N*_*e*_ between the replicates, which might be reflecting different selection histories possibly due to distinct genomic paths of adaptation.

### Recommendations for genome-wide data sets

Most of the methods proposed previously are not designed for genome-wide high density SNP data sets. However, the method of Jorde and Ryman (2007) was successfully used for genome-wide data by Foll *et al.* (2014). Reed *et al.* (2014) also used a similar approach to estimate *N*_*e*_ for whole genome data using sliding windows. We estimated *N*_*e*_ in windows with a fixed number of SNPs. Using windows of fixed lengths in base pairs, would affect the variance of the estimator (Fig. 4) but does not distort the mean. All these approaches however, do not account for the ruggedness of the recombination landscape and can lead to windows with different levels of linkage disequilibrium in them. To overcome this problem it would be possible to define windows based on recombination distance. Unfortunately, the lack of haplotype information in the Pool-Seq data makes it difficult to infer linkage disequilibrium. It is possible however to infer linkage information from the pooled sequence data using the software LDx (Feder *et al.* 2012). For model organisms, such as Drosophila readily available recombination maps can also be used as a proxy (Przeworski *et al.* 2001; Kulathinal *et al.* 2008; Fiston-Lavier *et al.* 2010). If only a single genome-wide *N*_*e*_ estimate is required, one can alternatively use a set of randomly distributed SNPs over the genome to obtain *N*_*e*_ estimate.

For our simulations, we considered only effective population sizes that are constant over time. Fluctuating *N*_*e*_ is a frequent phenomenon in natural populations and can be an important component of an experimental design. For example, in repeatedly bottlenecked populations, the smallest population size dominates the *N*_*e*_ estimate (Luikart *et al.* 1999; Charlesworth 2009). But even in strictly controlled populations the experimental regime can induce changes in *N*_*e*_. When the population changes in size the estimated *N*_*e*_ is generally interpreted as the harmonic mean of the effective population sizes over the generations (Wright 1938; Nei and Tajima 1981; Waples 1989). However, if time series allele frequency data is available such changes can be detected by estimating *N*_*e*_ from pairwise comparisons between consecutive time points.

To estimate *N*_*e*_ we used theoretical expectations of the variance in allele frequency after *t* generations of neutral evolution. All evolutionary forces (selection, demography, etc.) that lead to deviations from the neutral expectations will affect our estimate. Nevertheless, systematic forces that result in a localized reduction in *N*_*e*_, can be detected with the sliding window approach. The *Drosophila melanogaster* data set illustrates this point, i.e., the hypothesized region under selection exactly coincides with the region with reduced *N*_*e*_ (Orozco-terWengel *et al.* 2012; Tobler *et al.* 2014; Franssen *et al.* 2015).

### Using a small number of generations can lead to outlier estimates

In general, *N*_*e*_(*P*) has a lower bias but larger variance, especially when *t* is small. We observe outlier estimates among replicates at early generations (generation 5, Fig. 2 and S2-S4) for *N*_*e*_(*P*). The deviation from the true *N*_*e*_ is especially large when the sampling variance is large compared to the drift variance (*N*_*e*_ = 1000, P≤ 100 and c=50, Fig. S8,S9). As pointed out by Jorde and Ryman (2007) our weighting scheme leads to an increased variance but a smaller bias compared to other schemes. With this weighting design the signal for the *N*_*e*_ estimation is predominantly coming from the intermediate frequency variants. Having a large number of low frequency alleles thus has a similar effect as decreasing the number of SNPs.

To eliminate potential outliers and an inflated variance we recommend to increase the signal to noise ratio by pooling sufficient number of individuals. Using later generations or increasing the number of SNPs in the analysis also helps to avoid outlier estimates. When none of these strategies can be applied on the data, we suggest to use the genome-wide median taken across the estimates from individual windows because it is robust to outliers (Fig. 6). We also do not observe ouliers when *N*_*e*_(*P*) is applied on the genome-wide data set of *Drosophila melanogaster* (table 1).

## CONCLUSIONS

Effective population size an important parameter for describing evolutionary dynamics making its accurate estimation essential for population genetic studies. Several methods have been designed to estimate *N*_*e*_ for this purpose and their performance was comprehensively evaluated on simulated as well as real data (Gilbert and Whitlock 2015; Barker 2011; Serbezov *et al.* 2012; Baalsrud *et al.* 2014; Holleley *et al.* 2014). These studies mainly focused on genetic data collected from natural populations, which usually differs from experimental studies in terms of the census population size and sampling scheme. We designed a method that accurately infers the effective population size in genome-wide data from experimental populations sequenced in pools. Our approach improves temporal methods by explicitly correcting for two stages of sampling introduced by pooling and sequencing. Our results on simulated data confirm that methods that fail to properly account for the two stages of sampling inherent to Pool-Seq can lead to severely biased *N*_*e*_ estimates.

Pool-Seq data are often considered to be over-dispersed, i.e., displaying more variability than is predicted by the binomial sampling model (Yang *et al.* 2012). However, Zhu *et al.* (2012) and Futschik and SchlÖtterer (2010) validated that the error in allele frequency estimates is well approximated by binomial sampling given that a large enough number of individuals is pooled. We do not model over-dispersion in our correction term for pooling, nevertheless, it would be possible to introduce a parameter that accounts for the additional between-pool variation causing over-dispersion.

We illustrate the applicability of our method for estimating *N*_*e*_ from experimental data of Drosophila melanogaster and show that in combination with a recursive partitioning method we can infer patterns of local variation in *N*_*e*_ along the genome. Additionally, it is possible to calculate confidence intervals based on *χ*^2^ distribution (Nei and Tajima 1981) or alternatively apply non-parametric bootstrap approach and infer confidence limits based on sampling SNPs out of the original data set.

### Software availability

Our proposed estimators along with other methods from the literature are implemented within the R-package Nest. The package is currently available at https://github.com/ThomasTaus/Nest.

## ACKNOWLEDGMENTS

We thank Mads Fristrup Schou, Susanne U. Franssen and Neda Barghi for helpful comments on the *Nest* software package. AJ and TT are part of the Vienna Graduate School of Population Genetics which is funded by the Austrian Science Fund (FWF, W1225). CS is also supported by the European Research Council grant “ArchAdapt”, and TT is a recipient of a DOC Fellowship of the Austrian Academy of Sciences.

## List of Figures

1. Two-step sampling in experimental evolution with Drosophila 33
2. Performance of different methods to estimate *N*_*e*_ based on simulated data 34
3. Coefficient of variation of *N*_*e*_(*P*) for plan I for various parameter values 35
4. Effect of the number of SNPs used on the *N*_*e*_ estimates 36
5. Influence of the starting allele frequency distribution on *N*_*e*_(*P*) plan I estimator 37
6. Effect of linkage disequilibrium 38
7. Genome-wide *N*_*e*_ estimates from an E&R study in *D. melanogaster* 39

## List of Tables

1. Genome-wide mean and median *N*_*e*_ estimates from an E&R study in *D. melanogaster* 40

